# SLiMNet: a deep learning model to detect short linear motifs using protein large language model representations and paired inputs

**DOI:** 10.64898/2026.05.04.722713

**Authors:** Matthew C. McFee, Philip M. Kim

**Affiliations:** Department of Molecular Genetics, The University of Toronto, 1 King’s College, M5S3K3, Ontario, Canada; Department of Computer Science, The University of Toronto, 40 St George St, M5S2E4, Ontario, Canada; Donnelly Centre for Cellular and Biomolecular Research, The University of Toronto, 160 College St, M5S3E1, Ontario, Canada

**Keywords:** Deep learning, intrinsically disordered regions, short linear motifs, SLiM annotation

## Abstract

Short linear motifs (SLiMs) are short (3-15 amino acids in length) segments within intrinsically disordered regions (IDRs) that mediate transient protein–protein interactions as well as other functions such as stability and subcellular localization. Only a few thousand out of likely hundreds of thousands have been experimentally validated. SLiMs can be detected as conserved regions inside of IDRs using local alignments, though current approaches have limited sensitivity and specificity and are unable to functionally annotate their hits. Assigning function is hence a major outstanding issue in SLiM biology. Here we present SLiMNet, a deep learning model inspired by siamese networks and contrastive learning that predicts functional similarity in pairs of SLiMs. SLiMNet uses uses protein large language model embeddings and is trained on annotated sets of SLiMs. We show that it detects shared function in unseen, non-redundant motif pairs, and its scores correlate with experimental binding strengths from deep mutational scanning of cyclin-binding motifs. Using SLiMNet we provide repositories of putative SLiM pairs derived from annotated IDR regions for to help with hypothesis generation for the functional annotation of SLiMs. This includes an atlas generated from all-by-all scoring 16-mers from tiled IDRs from the DisProt database. We show that it captures a new nuclear localization motif recently added to MoMaP and a PRMT1 methylation motif in the literature. We also provided a repository of all IDRs scored with SLiMNet against against all MoMaP instances, and an atlas of potential functional pairs for 256 known orphan motifs (motifs with only a single known instance with essential function). Collectively, these atlases are useful resources for the SLiM biology community.

## Introduction

Short linear motifs (SLiMs) are 3-15 residue segments within intrinsically disordered regions that are associated with a variety of cellular functions, including transient protein-protein interactions, cellular localization, and influencing post-translational modifications [Davey et al., 2012]. Additionally, SLiMs are implicated in controlling protein localization, and stability. SLiMs have low levels of conservation, and often only a few amino acids in the motif may actually participate in an interaction [Van Roey et al., 2014]; hence SLiMs are difficult to identify. It is hypothesized that many thousands of functional SLiM classes are yet undiscovered [Bulavka et al., 2021].

Many different experimental techniques have been developed to find and determine short linear motifs and potential binding partners. One major technique is to generate libraries of peptides by tiling the IDRs of proteins associated with interacting with a protein of interest and then exposing these to bait proteins of interest or to use a functional readout to determine if the peptide is having some effect. For example, a pooled lentiviral system was to screen a library of 50000 peptides generated from IDRs and multiple inhibitors of the RWP1 cancer line [Nim et al., 2016]. An example of the former would be a recent study in which peptides were generated from IDRs in known cyclin interactors [Örd et al., 2025] and these peptides were assessed for binding ability to 11 different cyclins using the SIMBA system, [Subbanna et al., 2024] where peptides with high binding affinities will allow yeast to proliferate.

Methods for the computational detection of SLiMs has included developing local alignment algorithms to bypass the challenges of intrinsically disorder protein alignment (e.g. poor conservation) [Riley et al., 2023]. The goal of these methodologies is to improve IDR alignment to detect residue conservation with associated statistical confidences. For example, FaSTPACE [Kotb and Davey, 2024] performs pair-wise alignment of peptides to generated position-specific scoring matrices (PSSMs) and associated motifs. Similarly, the PairK [Halpin and Keating, 2024] method performs the alignment of k-mers extracted from an IDR believed to contain functional motif and aligns them with k-mers extracted from protein homologs to create multiple alignments of k-mers where conservation values can be computed and statistical significance assigned. These methods work on related principles and depend on some prior knowledge, PairK uses homolog sequences, and in the case of FaSTPACE you must provide the input set of peptides that you want to align to search for a motif. Furthermore, they only look for one potential feature for detecting/determining the existence of a SLiM-type.

Since SLiMs have significant functional importance, and have been an area of continued study for many years, several databases have been created to store information about experimentally validated, functional, SLiMs in proteins. The Eukaryotic Linear Motif (ELM) database [Kumar et al., 2023] is an important resource containing thousands of instances of validated motifs of various functions and is still being actively updated. More recently, MoMaP [Ambjørn et al., 2025] serves as a super-set of ELM with a large number of additional validated motif instances from a variety of curated sources. MobiDB [Piovesan et al., 2024] contains annotations for IDRs, included IDRs predicted by tools such as AlphaFold2 [Jumper et al., 2021] and includes other databases which annotate things such as disorder-to-order transitions in IDRs. Although relatively expansive, these databases are still thought to only cover a small minority of all SLiMs.

Large language models based on the Transformer architecture [Vaswani et al., 2017] have revolutionized natural language processing. By treating protein or nucleotide sequences as language, where each individual amino acid or nucleotide is represented as a token in the model architecture, protein large language models (pLLMs) have been trained on the large amount of available protein sequence data. The learned features in the representations of each amino acid have been shown to contain relevant information to predict many different biological targets [Schmirler et al., 2024]. In this work we utilize Evolutionary Scale Modeling 2 (ESM2) [Lin et al., 2022], a protein large language model trained on millions of sequences to generate informative features from protein sequences that can be used in conjunction with additional model modules to predict all-atom protein structures. ESM2 embeddings have been shown to be useful for many predictive tasks [Schmirler et al., 2024], even though the general usefulness of embedding spaces for downstream predictions has been a subject of debate. We were hypothesizing that pLLM embeddings have learned evolutionary signals which should make them more useful spaces to find and compare SLiMs than general sequence space.

## Results

### SLiMNet: a deep learning architecture that detects functional relationships between pairs k-mer inputs derived from intrinsically disorder regions

SLiMNet is a multi-layer perceptron (MLP), inspired by siamese neural networks [BROMLEY et al., 1993] and contrastive learning techniques [van den Oord et al., 2018] (Figure 1). In siamese networks, similarity between deep learning model outputs (typically pre-trained on a large dataset) are used for classification rather than classifying single inputs, allowing for new, rare categories of data to be classified against a few known examples. The SLiMNet architecture is as follows: the proteins containing the potential regions of interest, which may or may not contain a shared SLiM, are input into the ESM2 protein language model [Lin et al., 2022] (specifically the 650M parameter variant `esm2_t33_650M_UR50D`), and then the residues corresponding to the region of interest features are extracted from the final layer of the model and averaged to create a k-mer/motif level embedding. Any two k-mers of interest are then passed into the model and a similarity score, predicting whether or not there is a shared function between the k-mers (i.e. they are SLiMs of the same function) is returned. This technique also builds upon the technique used by SLiM detection methodologies like FaSTPACE [Kotb and Davey, 2024] and PairK [Halpin and Keating, 2024], algorithms which perform local alignments of IDR segments and then compute conservation metrics and associated p-values for statistical significance. Our model expands upon these models by allowing the similarity metric to be a learned function via deep learning instead of just alignment and conservation of residues. Furthermore, we utilize pLLM (protein large language model) embeddings which contain richer signals than sequence and have been shown to capture degrees of conservation in non-IDR regions [Yeung et al., 2023]. Finally, our network is novel in that we use the InfoNCE (defined in the Methods section) which trains the model to score related data points highest when considering a pool of possible matching candidates helping to alleviate against the inherent noisiness associated with SLiMs and IDRs.

**Fig. 1:**
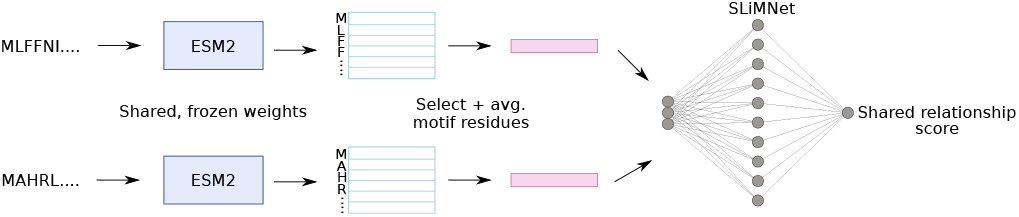
A high-level schematic representation illustrating the SLiMNet architecture. Our architecture is similar to siamese networks as the two parallel outputs from ESM2 are passed into SLiMNet, whose output is a score.

### SLiMNet can detect pairs with shared function vs. those without in a non-redundant hold-out test set derived from MoMaP

SLiMNet is trained on experimentally validated pairs of functionally related short linear motifs as positives, and negative pairs constructed by pairing motifs of unrelated functions to each other or real motifs against non-functional regular expression matches to any associated regular expression for the a given input motif. These sequences are all sourced from MoMaP (as of December 2024), a curated set of functionally annotated SLiMs maintained by Norman Davey’s group at the Institute of Cancer Research [Ambjørn et al., 2025]. This set is a super-set of the popular SLiM database, the Eukaryotic Linear Motif database [Kumar et al., 2023]. SLiMNet was initially tested on hold-out motifs with non-redundant functions (as annotated as `specificity_name` in MoMaP), in non-redundant sequences as determined by sequence similarity and clustering (consult Methods for clustering details). To reiterate, the model assigns scores to pairs consisting of two motifs of the same class, two motifs of different classes, or a motif and un-related regular expression match that is not functional. This set of points is constructed for each unique motif entry in the test set. Thus, if the model is performing well, it should apply high scores to the functionally related motif pairs and low scores to the pairs that are unrelated, which we quantified. When extracting the area under the receiver operating characteristic curve and the precision of the Precision-recall curve on the pairs constructed from these unseen SLiMs, SLiMNet generalizes well with an AUCROC (area under the curve of the receiver operating characteristic) and average precision of the Precision-recall curve of 0.81 and 0.71 respectively (Figure 2a,b). This suggests that SLiMNet is not simply memorizing certain sequences or memorizing specific classes of SLiM. Thus, we show that SLiMNet is generalizable to new motif types in novel sequences.

**Fig. 2:**
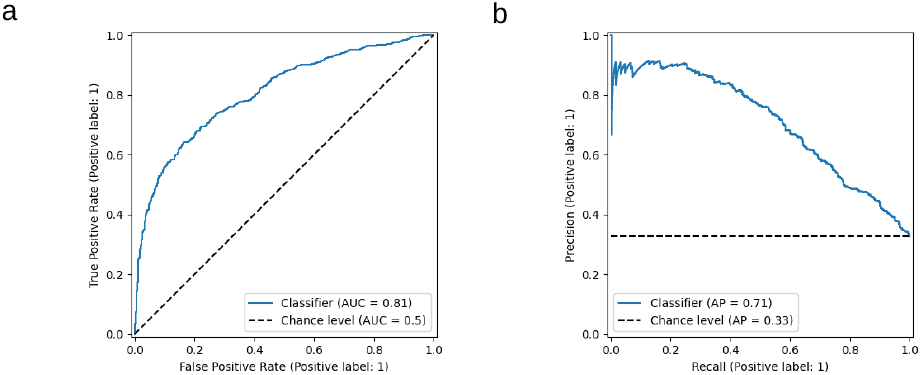
(a) Receiver operating characteristic curve of SLiMNet on hold-out non-redundant motif set. (b) Precision-recall curve of SLiMNet on hold-out non-redundant motif set.

### SLiMNet can detect novel cyclin binders that do not contain known cyclin binding motifs

Recently, a study [Örd et al., 2025] created an expansive library of 16 residue long k-mers constructed from the IDRs of proteins known to interact with 11 human cyclin variants. This generated library has tens of thousands of experimentally validated peptides with associated Systematic Intracellular Motif Binding Analysis (SIMBA) [Subbanna et al., 2024] binding strengths. Here, our model can differentiate reasonably well between positive pairs of novel cyclin binding peptides paired together versus negative pairs consisting of binders paired with non-binders. We quantify this performance with the AUCROC and the average precision of the precision-recall curve (Figure 3a,b). These read-outs will be particularly conservative because non-binders frequently contain regular expression matches to known cyclin binding motifs and may actually be functional SLiMs (but not cyclin binding motifs). Also, some of cyclin binding k-mers are very different from each other and all known cyclin binding motifs. Additionally, SLiMNet detects a functional relationship between input pairs when treating positive pairs as the novel cyclin binding regions from this study, paired with existing cyclin motifs contained in MoMaP and negative pairs as a known MoMaP cyclin binding motif and the k-mers determined not to bind in the screen [Örd et al., 2025] (Figure 4a, b). This case represents the use-case where a SLiMNet user would be searching for new instances of a known SLiM functional type by comparing against known instances. Interestingly, in the most challenging case, only selecting points where there is no regular expression match to the known cyclin motif, i.e. the most challenging possible binders from the screen as they are very diverse and have no discernible similarities to know binders, the signal reduces. This can be taken as the most challenging scenario for this data, but there is still a persistent signal in the precision-recall curve (Figure 4b).

**Fig. 3:**
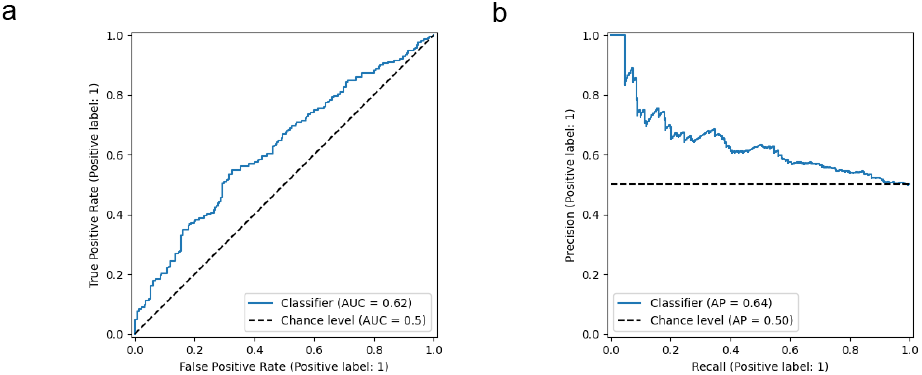
(a) Receiver operating characteristic curve of SLiMNet on hold-out non-redundant cyclin binding motif set. (b) Precision-recall curve of SLiMNet on hold-out non-redundant cyclin binding motif set.

**Fig. 4:**
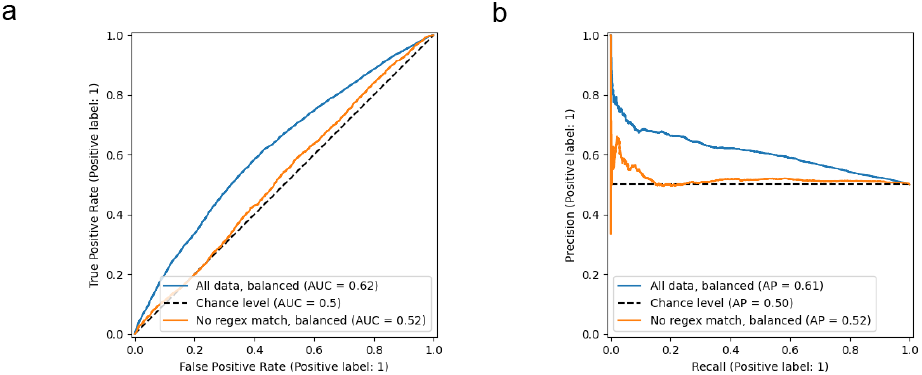
(a) Receiver operating characteristic curve of SLiMNet when pairing binders and non-binders against known cyclin binding motif instance inputs from MoMaP. Here pairs of novel binders and existing MoMaP instances of cyclins are positives, and negatives are a MoMaP binding instance, and a k-mer from the screen which was found to not bind any cyclin. Here we also include the scenario where we only include pairs that do not contain any known regex match to regular expressions associated with known cyclin binding SLiMs (orange curve). (b) Precision-recall curve of SLiMNet when pairing binders and non-binders against known cyclin binding motif instance inputs from MoMaP. Here positive and negative examples have been balanced to be in equal numbers. We also include the instance where we only include pairs that do not contain any known regex match to regular expressions associated with known cyclin binding SLiMs (orange curve).

### SLiMNet similarity scores of wild-type cyclin binding motifs and mutants correlate well with experimental binding data

In the cyclin screening paper [Örd et al., 2025], deep mutational scanning (DMS) was performed on a selection of cyclin binding motifs and SIMBA binding strength measurements were taken for all mutants. To assess how well our model can detect relative changes in binding affinity between the wild-type cyclin motif, and its mutants, the wild-type motif and each mutant are passed through the model as input pairs and the correlation of our scores to the to SIMBA scores were calculated. i.e. in our architecture one input would be the motif embedding extracted for the k-mer of interest in the wild-type sequence, the other input would be the mutant k-mer representation extracted from the protein sequence embedded with the appropriate amino acid modified in the input sequence. When selecting the top 100 highest scoring pairs from our model, there is a moderately strong correlation (Spearman of 0.44) with SIMBA binding values for many cyclins; we show cyclin A2 as an example in Figure 5a. However, when comparing the correlation to a position specific scoring matrix (PSSM) constructed using the SIMBA data from the paper, our model performs worse (Spearman *ρ* of 0.77 for the PSSM). Importantly, this PSSM will almost certainly perform better than any model that was not trained on highly redundant data, as the PSSM is constructed using deep mutational scanning data from all positions with associated binding strengths [Bandyopadhyay et al., 2020], thus it provides a rough upper performance bound for models of interest. To generalize this result, we plot the average Spearman *ρ* for the top 100 selections for each cyclin type between the SLiMNet and PSSM scores. It can be seen that although the mean of the SLiMNet scores appears lower (Figure 5b), there is no statistically significant difference in means as reported by a p-value of 0.09 from the Mann-Whitney U test. Moreover, the SLiMNet model scores correlated well with the PSSMs constructed using the binding preference data with a Pearson correlation of 0.66, *p* ≤ 0.05, indicating a significant positive linear correlation (Supplementary Figure A1). Because the PSSMs are constructed using the binding data, this is evidence that our model is learning information about binding strengths. Thus, although never trained on cyclin binding motifs, or any SLiM type from a protein known to bind a cyclin, our developed model seems to some extent be able to differentiate between mutations that increase binding affinity versus lower it. This further illustrates the power of SLiMNet for use with simulated motif deep mutational scanning studies.

**Fig. 5:**
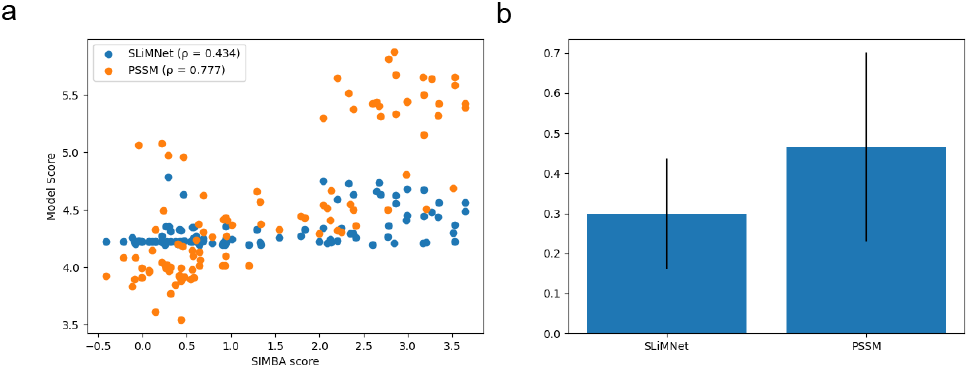
(a) Scatterplot comparing SLiMNet score to SIMBA value for cyclin A2. (b) Comparing mean Spearman correlations between our model and the position-specific scoring matrix. The difference is statistically significant under the Mann-Whitney U test (*p* = 0.09)

### SLiMNet can detect binding regions in IDRs within a protein of interest, the CAID Binding-IDR test set as a case study

As an additional validation of SLiMNet learning meaningful things about shared functional regions in IDRs, we applied SLiMNet to a dataset involving annotation of IDR regions being involved in binding. Here, we used the regions annotated in the Binding-IDR dataset from CAID3 [Mehdiabadi et al., 2025], again removing any SLiM in a sequence redundant to the Binding-IDR sequences from MoMaP before any training. This is a challenging test set of annotated regions within IDRs in proteins that are known to participate, or not participate in protein interactions. The model can differentiate between input pairs of two unique k-mers extracted from within annotated binding regions within a protein, from negative pairs consisting of a k-mer from a binding region, and a k-mer from a non-binding IDR in the same protein (Figure 6a,b). In other words, the model can determine if two unique portions of IDRs in a protein both participate in some interaction, from regions of IDRs in the same protein (the k-mers can be extracted from the same or different IDR regions) that do not bind anything. Here, the AUCROC is 0.78 and the AP is 0.78.

**Fig. 6:**
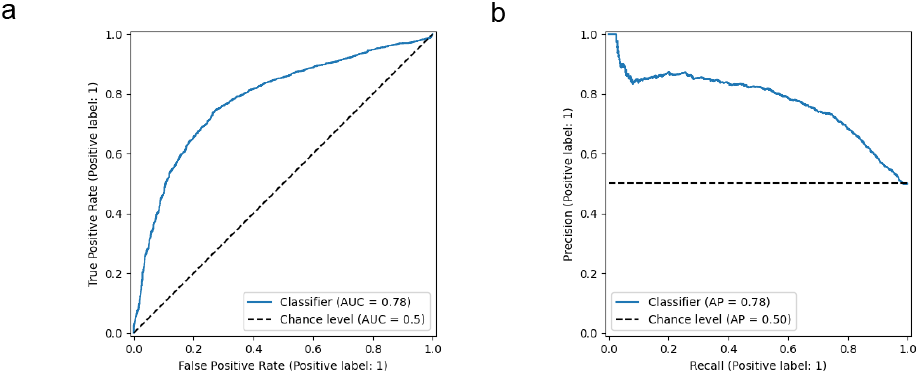
(a) The receiver operating characteristic curve for the Binding-IDR dataset. (b) The precision-recall curve for the Binding-IDR dataset.

However, it is important to note that the model does not classify well positive pairs consisting of k-mers that participate in interactions from different proteins. This indicates that our model is specific to motifs that bind the same thing, rather than general binding signal, which is enforced by our use of the InfoNCE loss. Motif pairs of the same binding type will score highly but those of different binding types will be negative points during training (see Methods). This result provides further support that SLiMNet learns meaningful functional similarities between extracted k-mer pairs. However, our model still performs well on the Binding-IDR set as the pairs are constructed within protein and there is shared binding signal at the protein level.

### Constructing an atlas of potential IDR regions with shared function in DisProt annotated human IDR regions as a resource for SLiM biologists

Next, we sought to create an atlas of pairs of potentially new functional regions within disorder regions, i.e. potential SLiM pairs as a resource for the community to explore and experimentally validate. The atlas consists of pairs of 16-mers, this length was selected to be in alignment with [Örd et al., 2025], constructed by tiling the disordered regions in human sequences obtained from DisProt [Aspromonte et al., 2024], a curated database containing annotations for disordered proteins and regions. The k-mers were compared in an all-by-all manner generated hundreds of millions of pairs of interest which were further filtered to the top 100,000 scoring pairs, as well as a subset of these with a Levenshtein difference of ≥ 10 to filter for 16-mers that are particularly different and likely representing novel SLiMs.

When selecting the top 100,000 pairs, it can be seen that the pairings consistently score highly with a right skew with an extended tail of exceptionally high scorers (Figure 7a). This indicates that there are many potential regions of interest for further biological study. Additionally, the Levenshtein distances [Berger et al., 2021] (the number of character edits to transform one sequence into the other) between the two k-mers is typically quite high indicating that the model is not simple scoring nearly identical regions of sequence as motifs of the same function (Figure 7c). Finally, when looking at a sequence logo, there is significant sequence diversity among the sequences in the pairs, although there does appear to be a large enrichment of glycines and lysines in the logo (Figure 7b) which is significantly different from the background (Figure 7d). Lysine and arginine enrichment are associated with nuclear localization signals [Lu et al., 2021], which is in alignment with our GO term enrichment analysis (Table 1) and the presented logo. However, there is likely some bias here since we tile the entire IDRs shifting a single residue at a time, so there are many related k-mers in the top 100000. Still, these serve as a reasonable starting point for further biological/experimental analysis.

**Fig. 7:**
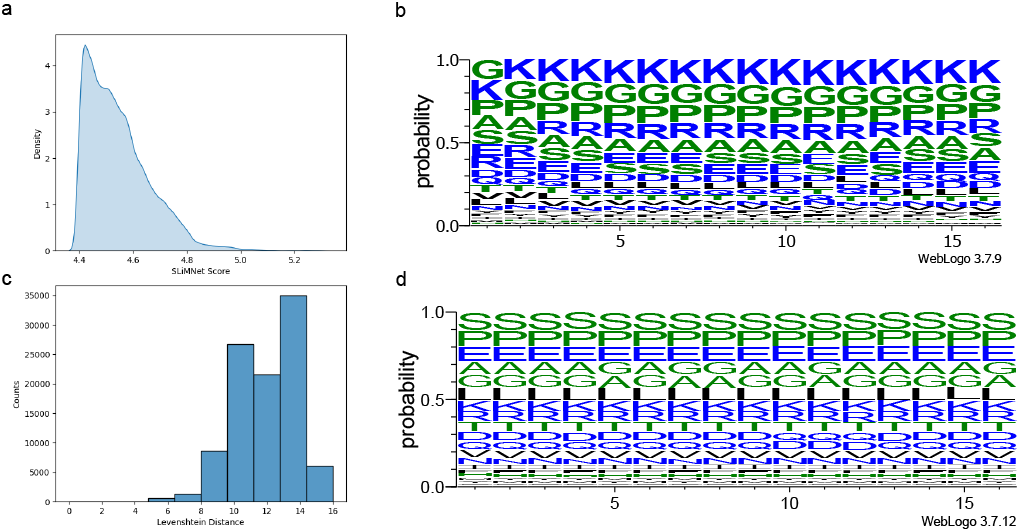
(a) Kernel density plot of the SLiMNet scores of the top 100000 scoring 16-mer pairs from the DisProt vs. Disprot library. (b) Histogram plot of Levenshtein distances between the 16-mers composing each unique pair. (c) Sequence logo (probability) of the unique sequences contained in the top 10000 scoring 16-mer pairs. (d) Sequence logo (probability) of the unique sequences contained in a random 100000 selected 16-mers from the whole Disprot versus Disprot k-mers library.

**Table 1.** ShinyGO Gene Ontology enrichment analysis for the Top 100K library. The top 10 enriched pathways are shown based on enrichment false discovery rate (FDR). This table was generated from initial analysis for the top 25 (Supplemental Table 1). Here we check enrichment of the overlap of pairs with STRING against the whole Top 100K library as the background.

| Enrichment FDR | nGenes | Pathway genes | Fold enrichment | Pathway |
| --- | --- | --- | --- | --- |
| 2.4E-06 | 16 | 1250 | 6.3 | Negative regulation of gene expression |
| 3.2E-05 | 7 | 163 | 20.2 | Heterochromatin formation |
| 4.9E-05 | 4 | 19 | 89.2 | Primary miRNA processing |
| 6.8E-05 | 7 | 199 | 16.8 | Negative regulation of gene expression, epigenetic |
| 7.7E-05 | 8 | 311 | 12.2 | Epigenetic regulation of gene expression |
| 7.7E-05 | 12 | 941 | 6.1 | Chromatin organization |
| 7.7E-05 | 20 | 2885 | 3.3 | Negative regulation of macromolecule metabolic process |
| 1.0E-04 | 5 | 72 | 31.2 | Regulatory ncRNA processing |
| 1.0E-04 | 18 | 2406 | 3.6 | Negative regulation of macromolecule biosynthetic process |
| 2.1E-05 | 14 | 1088 | 6.2 | RNA processing |

### Applications of the SLiMNet atlas of k-mers derived from human sequences in DisProt

As described above, we showed that the top 100000 scorers of our DisProt vs. DisProt atlas generated a diverse range of 16-mer pairs. we assessed the ability of the model to detect meaningful k-mer pairs in the atlas in two ways to illustrate the usefulness of our library.

More specifically, to assess whether or not the 16-mer pairs in the top 100000 selections (referred to as a Top 100K library in the remainder of this thesis) were in proteins with potentially associated functions, we considered interactions in the STRING database [Szklarczyk et al., 2023], in particular, the subset of STRING related to human proteins. STRING is a graph network where nodes are proteins and edges encode information about functional and regulatory relationships. Thus, we sought to determine if our model believes there are 16-mers with shared or related functions in proteins with known relationships in STRING. Interestingly, when checking our top scoring pairs, it can be seen that there is a marked increase in overlap of k-mer pairs coming from pairs with a known relationship in STRING in comparison to 100000 selected random pairs regardless of model score (Figure 8a, b). To assess if the model selects 16-mer pairs in the STRING related protein pairs with specific types of functions, the ShinyGO [Ge et al., 2019] tool was used to perform gene ontology (GO) term analysis, with all the sequences in the Top 100K acting as background. Here, we found a significant enrichment for many pathways associated with RNA and DNA regulation and processing (Table 1, expanded in Supplementary Table A1). Thus, it appears our model particularly can predict relationships between proteins associated with these pathways. This may, to some extent, be related to the consistent chemical properties or evolutionary signal of SLiMs associated with proteins localized to the nucleus.

**Fig. 8:**
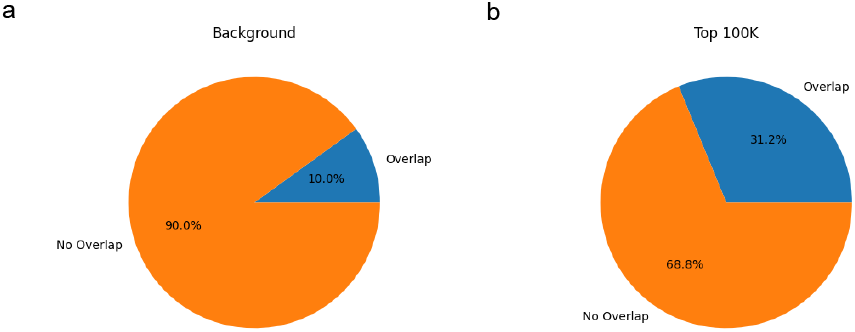
(a) Pairs of unique interacting proteins in 100K selected background pairs from all-by-all DisProt library. (b) The overlap with string for unique pairs in the Top 100K that contain no protein already in the MoMaP dataset. The difference in these two percentages of overlap were determined to be significant with the Chi-squared test *p* = 0.0.

To analyze this result further, the top scoring 16-mer pairs for any unique protein pairing in the Top 100K library that overlapped with STRING, and do not contain any protein existing in the version of MoMaP we trained on, were selected out. i.e. We examine only k-mer pairs in the Top 100K library that come from proteins that are paired in STRING and are not from sequences existing in MoMaP. We used this STRING filter to hone in on pairs in protein pairs with known functional relationships, as logically they may contain motifs of the same function. We then selected the top scoring k-mer pair for each unique pair of UniProt IDs. The top scorer of these pairs includes a k-mer from SERPINE1 mRNA-binding protein (UniProt ID Q8NC51) which matches a regular expression to `(GGRGG)|(.GRG.)` which is associated with MoMaP instances implicated in Protein Arginine Methyltransferase 1 (PRMT1) methylation. BLASTP [Altschul et al., 1990] returns no significant similarity between Q8NC51 and the proteins in MoMaP that contain this methylation site type. This hit in Q8NC51 appears in a pair with a 16-mer in the Heterogeneous nuclear ribonucleoprotein (hnRNP) that associates with mRNAs (UniProt ID P22626). Specifically, the 16-mer INFGDLGRPGRGGRGG in Q8NC51 scores highly with the 16-mer GNFGFGDSRGGGGNFG in P22626. Interestingly, there is no regular expression match for PRMT1 methylation in the k-mer from P22626, so searches with regular expression matches would fail to find this k-mer. To further validate this result, all MoMaP instances were paired against every generated 16-mers from DisProt and scored, with the Top 100K scoring pairs being selected. The 16-mer in Q8NC51 appears as a hit against the known MoMaP instance of a PRMT1 methylation site in Q01844, and many similar 16-mers in Q8NC51 score very highly with the instance of the methylation motif in Q6T6K0. An example of related 16-mers would be the 16-mer DLGRPGRGGRGGRGGR, which is shifted by only a few positions from the hit. This result further suggests that this pair is an instance of two PRMT1 methylation motifs.

The existence of methylation site motifs in two proteins associated with the ribosome and mRNA binding is logical due to the known functionality of PRMT1 methylation in regulating processes such as RNA processing [Sudhakar et al., 2023]. These methylation sites are known to regulate binding in the case of protein-protein and protein-nucleic acids [Blackwell and Ceman, 2012], including Q8NC51 [Ji and Ablimit, 2025]. In fact, this detected motif occurs in the region that has been proposed as a methylation site for PRMT1 which regulates mRNA binding [Lee et al., 2012]. Finally, hnRNP (P22626) is also known to contain sites which are methylated by PRMT1 [Sun, 2003, Huang et al., 2010, Li et al., 2021], and is associated with PRMT1 particles in general [Herrmann et al., 2005]. Thus, our model is proposing methylation sites in appropriate proteins (including the correct region in the case of Q8NC51), further proving the applicability of our model for motif discovery, including new instances of known functions.

MoMaP is a constantly evolving, curated dataset. Since the training and development of this model, new iterations with additional curated motif instances have been made available. Serving as a time-gated test set, similar to how models like AlphaFold3 [Abramson et al., 2024] are tested to see if the model performs well on new solved structures the model hasn’t possibly seen, we sought to see if the model recapitulated any new functional motif instances in a more recent version of MoMaP (version sourced in June 2025). Interestingly, in the top over-lapping scoring k-mer pairs that overlap with STRING, and that do not contain a sequence match to the trained on MoMaP instance, SLiMNet actually recapitulates a new annotated motif instance in the updated variant of MoMaP. In the latest variant of MoMaP, there is a new nuclear localization motif (AEKSKKKKEEEED, MoMaP ID IDII0000013572) in High mobility group protein B1 (UniProt ID P09429). This region appears as an exact match in our top scoring pairs in the correct protein (High mobility group protein B1). This protein is a chaperone protein in the nucleus, and is paired with other 16-mers in proteins such as Chromobox protein homolog 1 (P83916) which also associates with DNA and is also associated with DNA structure and regulation (Table 2). Thus, the model appears to be detecting novel nuclear localization motifs in proteins that are known to localize in the nucleus, as well as recapitulating a newly discovered motif of this type that is dissimilar to existing nuclear localization signals (no regular expression match to known nuclear localization motif regular expressions). Therefore, in two cases, we have shown that our library of potential related IDR regions recapitulates known functional SLiMs as supported by experimental literature.

**Table 2.**
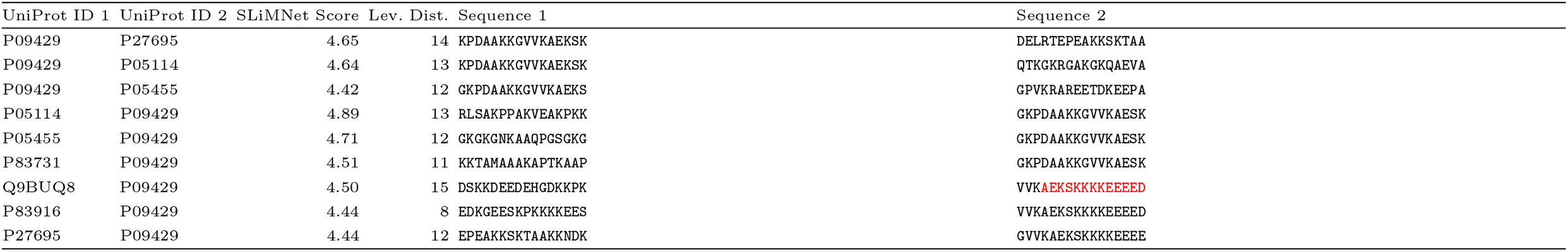
Similarity scores and Levenshtein distances between top scoring k-mer pairs from unique protein pairs in STRING, selecting only pairs similar to the nuclear localization signal motif (AEKSKKKKEEEED).

### Discovering potential functions of orphan SLiMs using against 16-mers constructed from the disordered regions of the entire human proteome

Orphan SLiMs are those that have been shown to be functional but for which only one instance of the motif exists. Here we focus on 252 orphan motifs that have been shown to be essential in recent work (publication forthcoming, data provided by Dr. Norman Davey). The embeddings of these SLiMs were then scored all-by-all against 16-mers generated by tiling across all regions annotated as curated/ideal from MobiDB [Piovesan et al., 2024] for the entire human proteome (accessed February 2026). The top 100000 scoring pairs were selected and characterized in the same way as in the previous section of this manuscript, again revealing diverse peptide pairs (Figure 9). Here we select the top scoring pairs from each unique pair of UniProt ID from MobiDB and the UniProt ID of each orphan. We present two high scoring pairs from the top two unique protein pairings as cases where our model is suggesting experimentally relevant pairings. First, we introduce the pair between DEADYVVPVEDNDENY and `YDQPW` in B-cell linker protein and SH2 domain-containing adapter protein D respectively. Both of these proteins act as adapter proteins connecting different proteins together. More specifically, SH2 domain-containing adapter protein D is known to contain functional `YXXP`motifs [Oda et al., 1997] which mediate protein interactions after phosphorylation. Thus, this serves as an experimentally supported, tangible related function in the B-cell linker protein, which is also associated with causing other proteins to undergo tyrosine phosphorylation [Baba et al., 2001], as well as having tyrosines within itself phosphorylated to facilitate function [Chiu, 2002]. In fact, `YXXP` motifs are associated with mediating SH2 domain binding [Liu et al., 2010] and are associated with SH2 binding in B-cell linker protein [Kabak et al., 2002], in particular modulating binding to proteins such as Phospholipase C, gamma 1. Here, our k-mer contains one of the known `YXXP` motifs phosphorylated by the protein Tyrosine-protein kinase SYK. This result is also further evidence that our model is producing pairs of high scoring k-mers that should be functional according to experimental evidence in the literature.

**Fig. 9:**
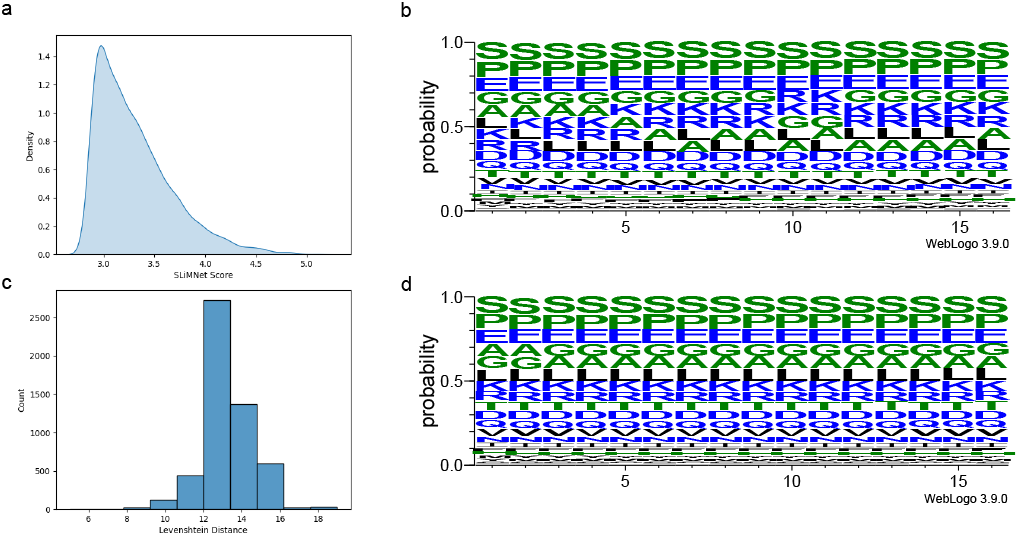
(a) Kernel density plot of the SLiMNet scores of the top 100000 scoring 16-mer from MobiDB and orphan motif pairs. (b) Histogram plot of Levenshtein distances between the two motifs composing each pair in the library. (c) Sequence logo (probability) of the unique sequences contained in the top 100000 MobiDB derived 16-mers from the library. (d) Background sequence logo (probability) of the unique sequences contained in all MobiDB derived 16-mers.

**Fig. 10:**
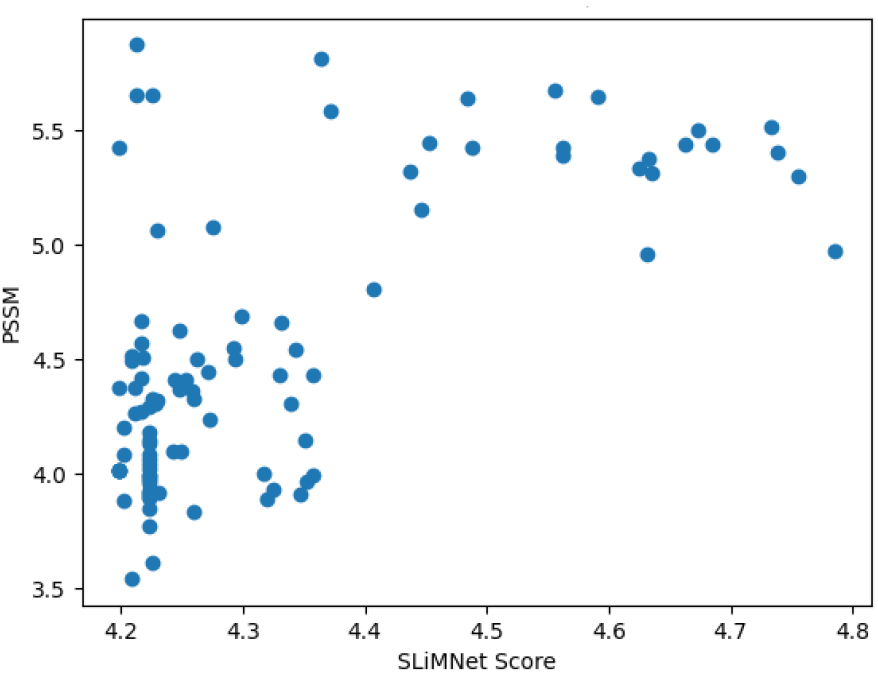
The top 100 highest scoring wild-type, mutant peptide pair scores correlate strongly with the maximum PSSM score for each motif from the available PSSMs generated in [Örd et al., 2025]. Pearson correlation of 0.66, with significant p-value. p ≤ 0.05.

Another case is Cyclic AMP-dependent transcription factor ATF-4 and Krueppel-like factor 4, with motifs DWMLEKMDLKEFDLDA and `LLDLDFILSN`, respectively. The motif in ATF-4 is in the N-terminal transcription activation domain (TAD) [Siang et al., 2025, Siang, 2023], and the orphan motif is near the 9-amino-acid TAD (9aaTAD) found in the literature [Piskacek et al., 2019], suggesting that this specific sub-region in ATF-4 could be responsible for TAD activity. Both proteins are known transcription factors associated with DNA binding, suggesting a potential shared role in mediating this function.

We also compared all MoMaP instances against the same MobiDB derived 16-mers presented in this portion of the manuscript for additional validation. We found that we can recapitulate the proposed novel phosphorylation motifs found with in our MoMaP versus DisProt library (Supplementary Table A2) such as Q8NC51 containing a PRMT1 phosphorylation motif. Furthermore, we identify new SH2 domain binding motifs in related immune system components. For example, Killer cell immunoglobulin-like receptor 3DL1 (P43629 UniProt ID) is proposed in the literature [Cheng et al., 2019] to bind SH2 domains such as that present in NKG2-A/NKG2-B type II integral membrane protein (IDII0000005433 in MoMaP).

Similarly, SLiMNet detects a SH2 binding motif again in UniProt ID P43629 and in MoMaP instance in Programmed cell death protein 1 (UniProt ID Q02242), another immune system associated protein. In this case, both proteins are known to bind SH2 domains to other proteins [Marasco et al., 2020, Cheng et al., 2018]. Thus, this result further supports the validity of our orphan motif predictions, and the ability of our model to detect binding domain motifs.

## Discussion

Here we introduce SLiMNet, a deep learning based-model that takes as input protein k-mer representations derived from amino acid embeddings of amino acids in IDRs extracted from the ESM2 protein large language model and predicts whether the input pair has shared functional relationships i.e. they are SLiMs of the same functional class. This work builds on recent works such as FaSTPACE and PairK which use local alignment tool of IDR regions to detect conserved residues participating in short linear motif function. By training on MoMaP, a curated dataset of experimentally validated and annotated short linear motifs, we show the functional power of SLiMNet. SLiMNet can detect shared function in pairs of unique instances of motifs of the same type in a non-redundant, hold-out test set of unseen motif functional types that are contained in sequences with low overall sequence identity (30% sequence identity clustering) to the sequences in the portion of MoMaP the model is trained upon.

Furthermore, we tested the ability of MoMaP on a recent library of cyclin binding motifs detected using SIMBA experimental binding analysis on a library of 16-mers generated in a tiling method from the IDRs of proteins associated with cyclin binding. To ensure no redundancy, we removed any sequence from MoMaP that clustered with the sequences used to generate the library, as well as removing any MoMaP instance whose specificity name (functional description) is associated with cyclin binding. We showed that SLiMNet can detect shared functional relationships in pairs constructed from these novel motifs, including the cyclin motifs with particularly unique compositions in comparison to known SLiMs involved in cyclin binding. Interestingly, when pairs of cyclin binding motifs and their associated mutants from deep mutational scanning experiments are used as input pairs to our model, we find that our model scores correlate well with the SIMBA binding scores for each cyclin type when selecting for the top 100 highest scoring pairs according to SLiMNet. Although performing worse than PSSM generated scores in many cases, the PSSM scores act as an upper bound to model performance as they are directly generated using experimental SIMBA data from the completed DMS data, and act as an upper bound of performance. Statistical analysis across all correlations reveals our model performs similarly to PSSMs when looking at the top 100 scoring pairs of wild-type and mutant according to our model scores. This result indicates that our model is learning about function and binding affinity.

We also show that our model has predictive signal in the Binding-IDR dataset, from CAID3, a challenging set for testing whether or not regions of IDRs within a protein participate in transient interactions. Here, our model can differentiate k-mer pairs generated from unique binding regions as being both involved in binding from pairs of k-mers where one is a binder and one is a non-binder. This result is at a per protein level.

The final contributions of this work include several atlases of pairs of 16-mers extracted by tiling the IDRs of human proteins as contained in DisProt or MobiDB. We subset these into smaller atlases of 100000 (a Top 100K library) of the top scoring pairs across the millions of computed values which show high scores, which are still diverse as indicated by large edit distances (as quantified by Levenshtein distances) between the k-mers in the pairs. These are a promising initial point for biologists to explore, but we provide access to all computed pairs (all-by-all). In the case of 16-mers constructed using DisProt and scored all-by-all, our analysis of the Top 100K selections from the library, we find two promising cases for further experimental validation. We find a relevant new methylation site in SERPINE1 mRNA-binding protein 1, in the top scorers, that also appears in the Top 100K scoring pairs when all the 16-mers generated from DisProt are scored against the known MoMaP instances from the iteration of MoMaP used to construct the model. We also show that our model also predicts a new motif in a more recent version of MoMaP that the model could not have seen as a functional nuclear localization motif. Thus, SLiMNet can rediscover unseen/annotated motif instances that have now been experimentally validated as real and functional and deposited in MoMaP. Additionally, we provide evidence for new instances of orphaned motifs by doing an all-by-all comparison of known orphans to 16-mers derived from the entire human proteome IDRs as annotated by MobiDB. We suggest functions and new instances for two key orphan motifs and is paired with a known `YXXP` motif in another protein, with experimental support in the literature. The other orphan motif is likely associated with transcription activation as it has been recently been experimentally shown in the literature, the other region it pairs with is also in a region associated with transcription activation, and thus the paired k-mer is likely functional in transcription activation. Re-scoring all MoMaP motifs against this MobiDB derived library for the entire proteome re-capitulates results from our MoMaP versus DisProt library, while providing evidence for new SH2-binding motifs in immune-system related proteins. These results cumulatively show that our model can be used to discover novel motif instances.

Although a promising new model, future work can be done to strengthen the value of SLiMNet. First and foremost, promising pairs from the atlas should be further analyzed by biologists with the most promising instances to be validated experimentally. It would also be valuable to benchmark the model on new motif instances as they become available.

## Methods

### Training details

The model is a two layer multi-layer perceptron that features layer normalization and dropout for regularization. The input data contains 2560 features, and the hidden layers contain 640, and 320 hidden neurons, respectively. These values correspond to roughly 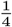 and 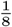 the number of input features. The final weights map to a single similarity score which is passed to the InfoNCE loss for training. The model was trained until convergence using the InfoNCE loss and the AdamW optimizer with a learning rate of 1*e* − 5, with a batch size of 64, and a dropout rate of 0.25. Please refer to the included model as needed.

### The InfoNCE loss

The InfoNCE loss [van den Oord et al., 2018] is a contrastive learning objective where given a data point and a similarity metric, the InfoNCE should provide the highest similarity to another data point of the same function and low similarities to unrelated data points. Effectively, the loss is a cross-entropy loss where the highest probability should be assigned to the pair of inputs which are most similar. Here we adapt the notation presented in [Lu et al., 2022]

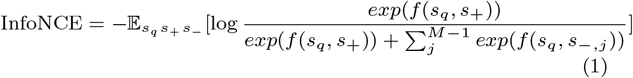

Here *s*_*q*_ is the query input motif, *s*_+_ is a different motif from the same functional class and *s*_−_ is a negative point consisting of a potentially a motif of another function with no relation to the query, or a non-functional protein segment that contains a regular expression match to the query. M is the total number of points the query is being compared to, here there is 1 + example, and M-1 negatives. The function *f* () is the similarity function to determine the relationship between the points (SLiMNet in this presented work), motifs of the same function should have large positive assigned scores. Thus, this function represents the probability that the query and its positive match are assigned a high probability against a negative background set after applying the softmax function. The expectation is approximated via a batch of input points during training being averaged.

### dataset splitting and redundancy reduction

For the MoMaP training validation, and test sets, the sets were created to approximate a 80/10/10% split where there was no overlap in specificity name (functional annotation) between the motifs included in any set and no cluster overlap at 30% sequence identity when clustering with the MMSeqs2 algorithm [Steinegger and Söding, 2017]. Any MoMaP instance with a functional annotation associated with cyclin binding was removed from the training data. Additionally, all the sequences included in the MoMaP database were clustered against the sequences from which the cyclin and Binding-IDR datasets were generated and redundant sequences in the training, validation, and test sets derived from MoMaP were removed. Any MoMaP instance in a sequence that clustered at 30% identity to any of these sequences was also removed to prevent redundancy in the training and test data sets.

### Gene enrichment analysis details

Here we apply the ShinyGO tool [Ge et al., 2019] for GO term enrichment analysis, particularly biological process GO terms. The settings we used for this analysis are as follows: the false discovery rate (FDR) is 0.05, 25 pathways are displayed, the minimum pathway size was 2, the maximum pathway size was 5000 and all redundancy was removed. Here are overlapping pairs that are in STRING and neither member of the pair is in the MoMaP instance we used for training. The background are all the proteins contained in the Top 100K pairs library in the case of the Disprot Vs. Disport library.

## Author contributions statement

The concept of the paper was developed by M.C.M. Data generation, curation, and analysis was performed by M.C.M. Model design and implementation was performed by M.C.M. Writing of this manuscript was performed by M.C.M. with support from P.M.K. Supervision and funding acquisition was by P.M.K.

## Acknowledgments

We would like to thank the Digital Research Alliance of Canada for providing computing resources used to generate the data of this paper. We would like to thank Norman Davey for his support, insightful feedback, and with providing the MoMaP and orphan motif data for developing SLiMNet.

## Code and data availability

All code and data (including all raw data and the entire top 100000 selections for Top 100K libraries) are available upon request to the corresponding author.

## Supplemental Figures

## Supplemental Tables

**Table A1:**
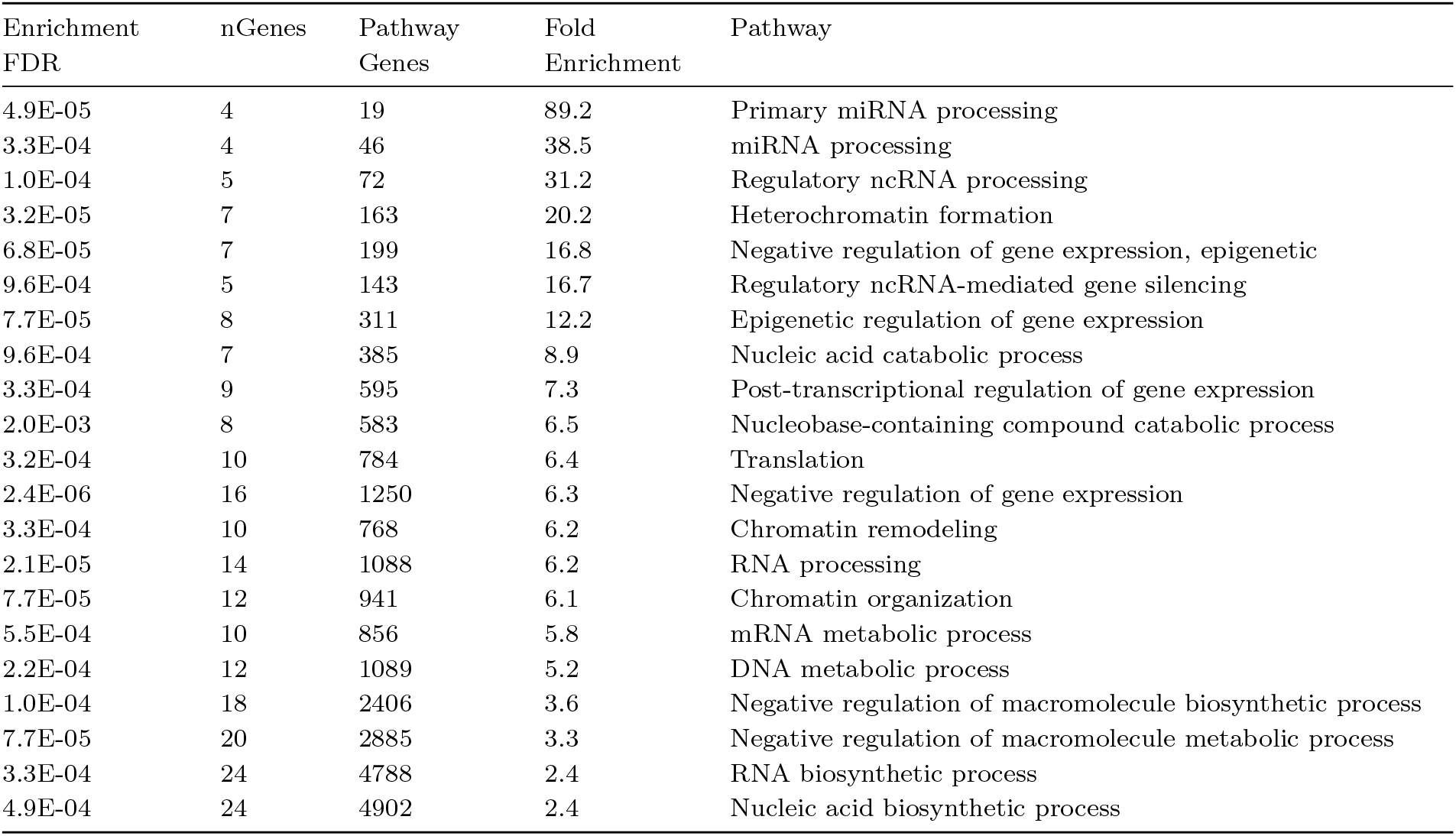
Extended ShinyGO GO enrichment analysis for the Top 100K library.

**Table A2.**
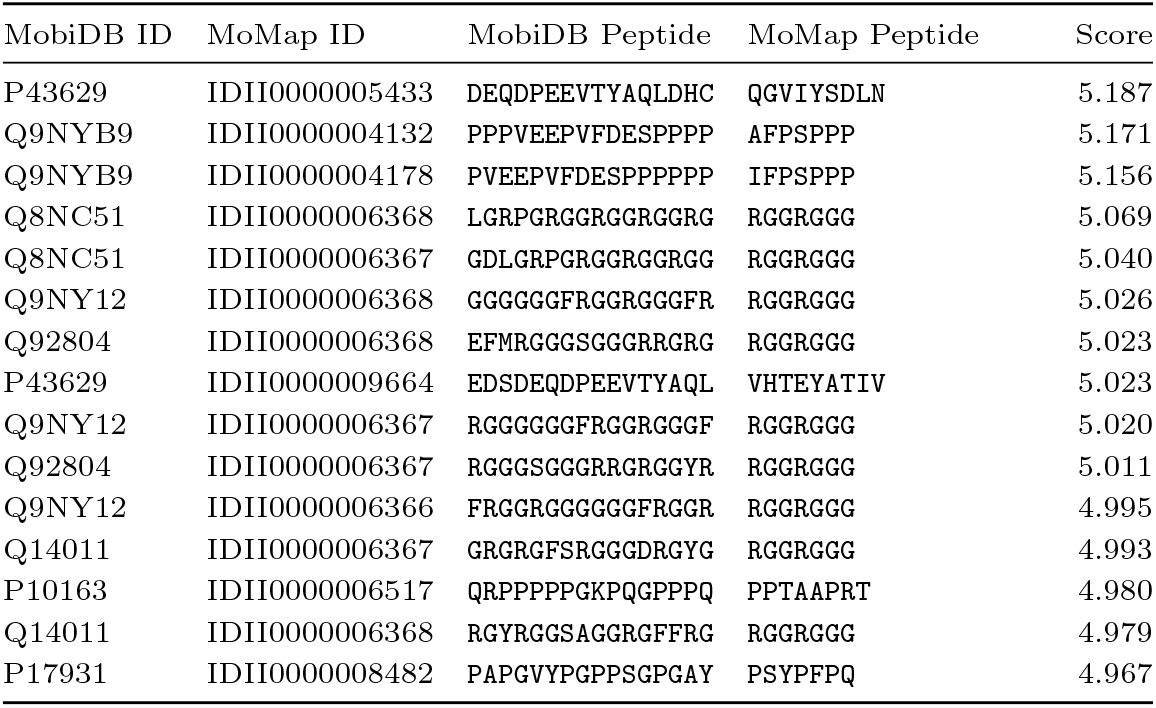
Fifteen top scoring peptide similarity matches between unique MobiDB UniProt IDs that are not present in the version of MoMaP used to train SLiMNet and MoMaP instances. This table is generated by filtering the entire top 100000 scoring pairs from the entire MobiDB versus MoMaP library for top scoring pair of all unique pairings of MobiDB UniProt ID and MoMaP ID.

