## supplemental for "SLiMNet: a deep learning model to detect short linear motifs using protein large language model representations and paired inputs"

### Supplemental Figures

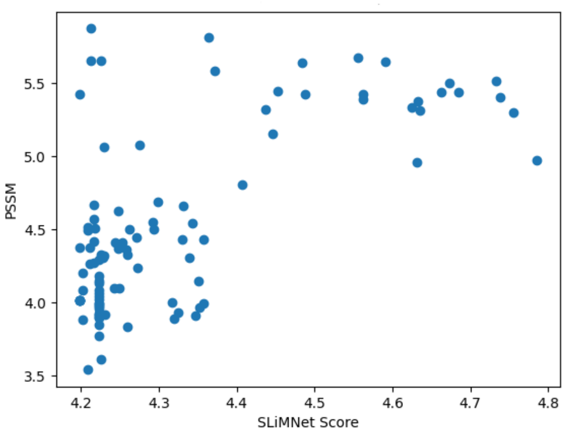


Fig. 10: The top 100 highest scoring wild-type, mutant peptide pair scores correlate strongly with the maximum PSSM score for each motif from the available PSSMs generated in [Örd et al., 2025]. Pearson correlation of 0.66, with significant p-value. $p \leq 0.05$.

### Supplemental Tables

Table A1: Extended ShinyGO GO enrichment analysis for the Top 100K library.

| Enrichment FDR | nGenes | Pathway Genes | Fold Enrichment | Pathway |
|---|---|---|---|---|
| 4.9E-05 | 4 | 19 | 89.2 | Primary miRNA processing |
| 3.3E-04 | 4 | 46 | 38.5 | miRNA processing |
| 1.0E-04 | 5 | 72 | 31.2 | Regulatory ncRNA processing |
| 3.2E-05 | 7 | 163 | 20.2 | Heterochromatin formation |
| 6.8E-05 | 7 | 199 | 16.8 | Negative regulation of gene expression, epigenetic |
| 9.6E-04 | 5 | 143 | 16.7 | Regulatory ncRNA-mediated gene silencing |
| 7.7E-05 | 8 | 311 | 12.2 | Epigenetic regulation of gene expression |
| 9.6E-04 | 7 | 385 | 8.9 | Nucleic acid catabolic process |
| 3.3E-04 | 9 | 595 | 7.3 | Post-transcriptional regulation of gene expression |
| 2.0E-03 | 8 | 583 | 6.5 | Nucleobase-containing compound catabolic process |
| 3.2E-04 | 10 | 784 | 6.4 | Translation |
| 2.4E-06 | 16 | 1250 | 6.3 | Negative regulation of gene expression |
| 3.3E-04 | 10 | 768 | 6.2 | Chromatin remodeling |
| 2.1E-05 | 14 | 1088 | 6.2 | RNA processing |
| 7.7E-05 | 12 | 941 | 6.1 | Chromatin organization |
| 5.5E-04 | 10 | 856 | 5.8 | mRNA metabolic process |
| 2.2E-04 | 12 | 1089 | 5.2 | DNA metabolic process |
| 1.0E-04 | 18 | 2406 | 3.6 | Negative regulation of macromolecule biosynthetic process |
| 7.7E-05 | 20 | 2885 | 3.3 | Negative regulation of macromolecule metabolic process |
| 3.3E-04 | 24 | 4788 | 2.4 | RNA biosynthetic process |
| 4.9E-04 | 24 | 4902 | 2.4 | Nucleic acid biosynthetic process |

**Table A2.** Fifteen top scoring peptide similarity matches between unique MobiDB UniProt IDs that are not present in the version of MoMaP used to train SLiMNet and MoMaP instances. This table is generated by filtering the entire top 100000 scoring pairs from the entire MobiDB versus MoMaP library for top scoring pair of all unique pairings of MobiDB UniProt ID and MoMaP ID.

| MobiDB ID | MoMap ID | MobiDB Peptide | MoMap Peptide | Score |
|---|---|---|---|---|
| P43629 | IDII0000005433 | DEQDPEEVTYAQLDHC | QGVIYSDLN | 5.187 |
| Q9NYB9 | IDII0000004132 | PPPVEEPVFDESPPPP | AFPSPPP | 5.171 |
| Q9NYB9 | IDII0000004178 | PVEEPVFDESPPPPPP | IFPSPPP | 5.156 |
| Q8NC51 | IDII0000006368 | LGRPGRGGRGGRGGRG | RGGRGGG | 5.069 |
| Q8NC51 | IDII0000006367 | GDLGRPGRGGRGGRGG | RGGRGGG | 5.040 |
| Q9NY12 | IDII0000006368 | GGGGGGFRGGRGGGFR | RGGRGGG | 5.026 |
| Q92804 | IDII0000006368 | EFMRGGGSGGGRRGRG | RGGRGGG | 5.023 |
| P43629 | IDII0000009664 | EDSDEQDPEEVTYAQL | VHTEYATIV | 5.023 |
| Q9NY12 | IDII0000006367 | RGGGGGGFRGGRGGGF | RGGRGGG | 5.020 |
| Q92804 | IDII0000006367 | RGGGSGGGRRGRGGYR | RGGRGGG | 5.011 |
| Q9NY12 | IDII0000006366 | FRGGRGGGGGGFRGGR | RGGRGGG | 4.995 |
| Q14011 | IDII0000006367 | GRGRGFSRGGGDRGYG | RGGRGGG | 4.993 |
| P10163 | IDII0000006517 | QRPPPPPGKPQGPPPQ | PPTAAPRT | 4.980 |
| Q14011 | IDII0000006368 | RGYRGGSAGGRGFFRG | RGGRGGG | 4.979 |
| P17931 | IDII0000008482 | PAPGVYPGPPSGPGAY | PSYPFPQ | 4.967 |
